# Quantifying Behavioral Structure and Persistence in Open-Field Assays Using Entropy and Spectral Metrics

**DOI:** 10.64898/2026.02.20.707059

**Authors:** Sihak Lee, Daniel J. Choi, Soonwook Choi, Zhanyan Fu

## Abstract

Traditional open-field assays quantify rodent behavior using low-dimensional scalar metrics that often overlook dynamical and temporally dependent aspects of spontaneous behavior. Here, we establish a high-throughput framework that integrates multi-camera 3D pose estimation from AVATAR-3D (1) with unsupervised autoregressive hidden Markov modeling using Keypoint-MoSeq (2–4) to transform recordings into time sequences of discrete, recurring behavioral units termed syllables. Drawing inspiration from information theory (5) and spectral analysis of Markov systems (6,7), we compute Shannon entropy and the second-largest eigenvalue (Eigen_2_) of the syllable transition matrix. Entropy and Eigen_2_ offer complementary lenses through which spontaneous behavior can be understood—one capturing dispersion of behavioral expression, the other reflecting temporal persistence and mixing dynamics. Together, these metrics quantify the organization of behavior as a stochastic dynamical system rather than a collection of isolated actions.

## OVERVIEW

### Introduction

Traditional openfield assays have relied on low-dimensional scalar metrics such as average velocity, total distance travelled, or time spent in the center of the arena. While these measures provide insights into subjects’ locomotor patterns and preferences, they overlook potentially rich temporal intricacies of behavior. With the advent of machine learning models that can serve as hands-free unsupervised magnifiers, a more complex yet intriguing type of analysis has become possible.

By combining two novel technologies, AVATAR 3D and Keypoint Moseq (KPMS), a unique high-throughput pipeline digests openfield recordings in a dynamical, data-driven manner. Supplemented by additional analysis separate from either imported frameworks, providing a complementary and more structured characterization of spontaneous behavior relative to traditional scalar measures.

AVATAR 3D provides necessary hardware and software necessary to record freely moving mice in its arena and produce 3D coordinate data of landmarks. At a high level, this is accomplished by markerless pose estimation from each of its angles and then triangulation. The coordinate data is restructured to be fed into KPMS, where an unsupervised modelling framework segments the continuous coordinate data into discrete, recurring units called “syllables.” In other words, rather than relying on predisposed top-down knowledge of frequent or expected mouse behaviors, the model is able to identify repeated patterns that emerge bottom-up directly from the data.

Prevalent mouse behaviors can be retroactively labelled. What we have is a temporally organized sequence of syllables that can be analyzed across cohorts. KPMS provides a suite of informative analyses that can be applied to syllable-annotated behavioral sequences. For example, syllable frequencies and transition counts can be quantified and compared across cohorts, and the most prevalent state transitions can be identified and ranked. While these measures offer valuable insight into specific behavioral motifs, they remain largely state-level and descriptive. As analyses become increasingly granular—such as aggregating selected syllables that resemble a particular behavior and comparing their combined frequency across groups—interpretation can become dependent on subjective grouping decisions. Although such comparisons are informative, they do not provide a concise summary of the overall structural organization of behavior. To address this limitation, we sought metrics capable of characterizing the behavioral system as a whole, rather than focusing solely on individual syllables or selected subsets.

Hence, we introduced eigenvalues and Shannon’s entropy into a modified version of KPMS’s output. From the temporally ordered syllable sequence, we derived an (N+1) X (N+1) transition probability matrix, where S={syllables (0,…,N) ∪ other}and each entry represents the probability of transitioning from one syllable to another. The matrix was constructed to be row-stochastic, ensuring that the probabilities of all possible future states conditioned on a given state sum to one.

This representation transforms the behavioral sequence into a discrete stochastic dynamical system. From this structure, Shannon’s entropy provides a measure of how behavioral states are distributed across time, while the second largest eigenvalue of the transition matrix captures the rate at which behavioral sequences mix and persist (5-7). Together, these complementary quantities offer a succinct characterization of both the distributional and temporal organization of spontaneous behavior.

### Methods

Whereas KPMS proposes using tools such as SLEAP or Deep Lab Cut (DLC) for recording and coordinate acquisition, AVATAR 3D was used instead for higher accuracy (1). For most single depth camera set-ups, as in most DLC projects, coordinate acquisition relies on a wider confidence interval; for some frames where certain landmarks become obscured, the coordinates acquired inherently possess lower accuracy. With AVATAR 3D’s 20cm x 20cm arena and 5 camera setup that encompasses every side of the cube except the top, estimating coordinates for hidden body parts no longer is an issue. The cameras acquire data in sync at 30 frames per second.

Pose estimation was performed using a pre-trained deep neural network targeting to detect 9 anatomical landmarks in mice: nose, neck, body center, four limbs, anus, and tail tip. Tail Tip was later discarded as it was deemed to contribute more to noise than meaningful data in subsequent modelling. For each frame, two-dimensional landmark detections from all cameras were integrated through automatic multi-view triangulation to reconstruct three-dimensional coordinates while minimizing occlusion-related ambiguity.

AVATAR 3D’s platform also included the ability to process output coordinate data as a stick-figure animation. The animations for each respective recording served as visual verification of the acquisition’s accuracy and continuity.

The final output of this stage consisted of temporally ordered 3D coordinate trajectories for each retained landmark.

Raw 3D coordinate data from AVATAR 3D were restructured using a custom data loader to match the input format required by KPMS. Coordinates were organized into time-indexed arrays with consistent ordering of landmarks. All confidence values were assumed to be 1.

KPMS framework were initialized using a project-level configuration file (YAML) based on recommended defaults for mouse pose modeling, which was adopted and modified as appropriate The configuration specified: the full set of tracked body parts and the subset used for modeling (excluding the tail tip), a skeletal graph denoting bodypart connectivity, anterior/posterior reference points for orientation, confidence handling parameters (e.g., confidence threshold and pseudocount), whitening, and priors/hyperparameters governing observation noise, autoregressive dynamics, and state transition structure.

To reduce redundancy among correlated keypoint trajectories and improve computational efficiency during model inference, dimensionality reduction was performed using Principal Component Analysis (PCA).

The original pose representation consisted of three-dimensional coordinates (x, y, z) for each of eight retained anatomical landmarks per frame. These coordinates were concatenated into a high-dimensional pose vector at each time point. PCA was applied across the full dataset to identify orthogonal components capturing the dominant modes of postural variation.

The number of retained principal components was capped at 10, or fewer if a smaller number of components explained ≥ 90% of the total variance. This reduced representation preserved the majority of postural variance while eliminating redundant dimensions and stabilizing downstream behavioral segmentation.

The resulting lower-dimensional trajectories served as input to the Keypoint-MoSeq modeling framework.

Behavioral segmentation was performed using the KPMS autoregressive hidden Markov model (AR-HMM) framework, which infers a discrete sequence of behavioral states (“syllables”) from the reduced-dimensional pose trajectories (2,3).

Model fitting proceeded in two stages. First, the model was trained in an autoregressive-only regime (AR-only) for an initialization phase of 200 iterations. This warm-start procedure stabilizes inference of autoregressive dynamics before incorporating additional non-autoregressive structure. The resulting checkpoint was then used to initialize full model training that included both AR and non-AR components, which was continued for an additional 500 iterations.

To maintain the intended syllable time scale (i.e., to control state persistence/dwell times), we adjusted the transition hyperparameter kappa prior to the second training stage, following KPMS guidance for tuning syllable duration. Unless otherwise noted, all other hyperparameters were retained from the configuration defaults.

The final trained model produced, for each recording, a time-indexed sequence of inferred syllable labels and associated posterior quantities used for downstream analyses.

The syllable-annotated sequences naturally can be used to measure the syllable count, frequency, as well as the transitional counts and probability. These metrics were plotted to show a comparison between cohorts.

For additional analysis, a syllable transition probability matrix P∈R(N+1)×(N+1) was constructed over the state space:

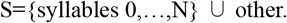

Each matrix was constructed to be row-stochastic, meaning:

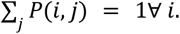

In cases where a syllable was not observed within a segment, smoothing was applied to preserve row-stochasticity and enable stable spectral analysis (see below).

Global entropy was computed from the stationary distribution of syllable occupancy:

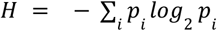

where *p*_*i*_ represents the empirical probability of syllable i within the recording or segment.

Local entropy was computed for individual syllables based on their outgoing transition distributions:

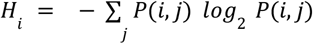

This measure captures the dispersion of next-state transitions conditional on being in state i.

The eigenvalues of the transition matrix P were computed via standard eigendecomposition.

Because P is row-stochastic and a square matrix, its largest eigenvalue is 1. The second largest eigenvalue in magnitude (Eigen_2_) was extracted and used as a standard metric of temporal persistence and mixing dynamics.

Values closer to 1 indicate slower mixing and increased persistence in behavioral transitions, whereas smaller values indicate more rapid mixing and reduced temporal dependency.

## RESULTS

### Characterizing the Spatiotemporal Structure and Stability of Spontaneous Behavioral Repertoires in Wild-Type Mice

In order to demonstrate the full capability of our dynamical behavior pipeline, we applied the framework to 30-minute openfield recordings of twelve wild-type C57BL/6J (WT) mice. These animals were originally collected as a set of twenty-four – twelve of which were unused complimentary recordings for the Mutant SETD1A genotype; in this section, the WT mice were exclusively used to illustrate the structure, interpretability and multi-scale outputs of the pipeline independent of experimental manipulation.

Figure 1A represents what was much already discussed in the methods section. Mice recordings hosted in AVATAR3D are processed into 3D pose trajectories at 30 frames per second, and they are introduced into KPMS where they are subsequently reduced into simpler space via principal component analysis and segmented into discrete behavioral syllables. This transformation gives rise to an annotated version of the original recording file that illuminates what syllable each frame belongs to – according to modeling parameters. The standard outputs generated by KPMS, which include syllable frequency as well as transition counts, are complemented by custom transition-based and information-theoretic analyses. Together, these steps establish a unified dynamical representation of spontaneous behavior that serves as the methodological foundation for subsequent pharmacological, environmental, and genetic comparisons.

**Figure 1.**
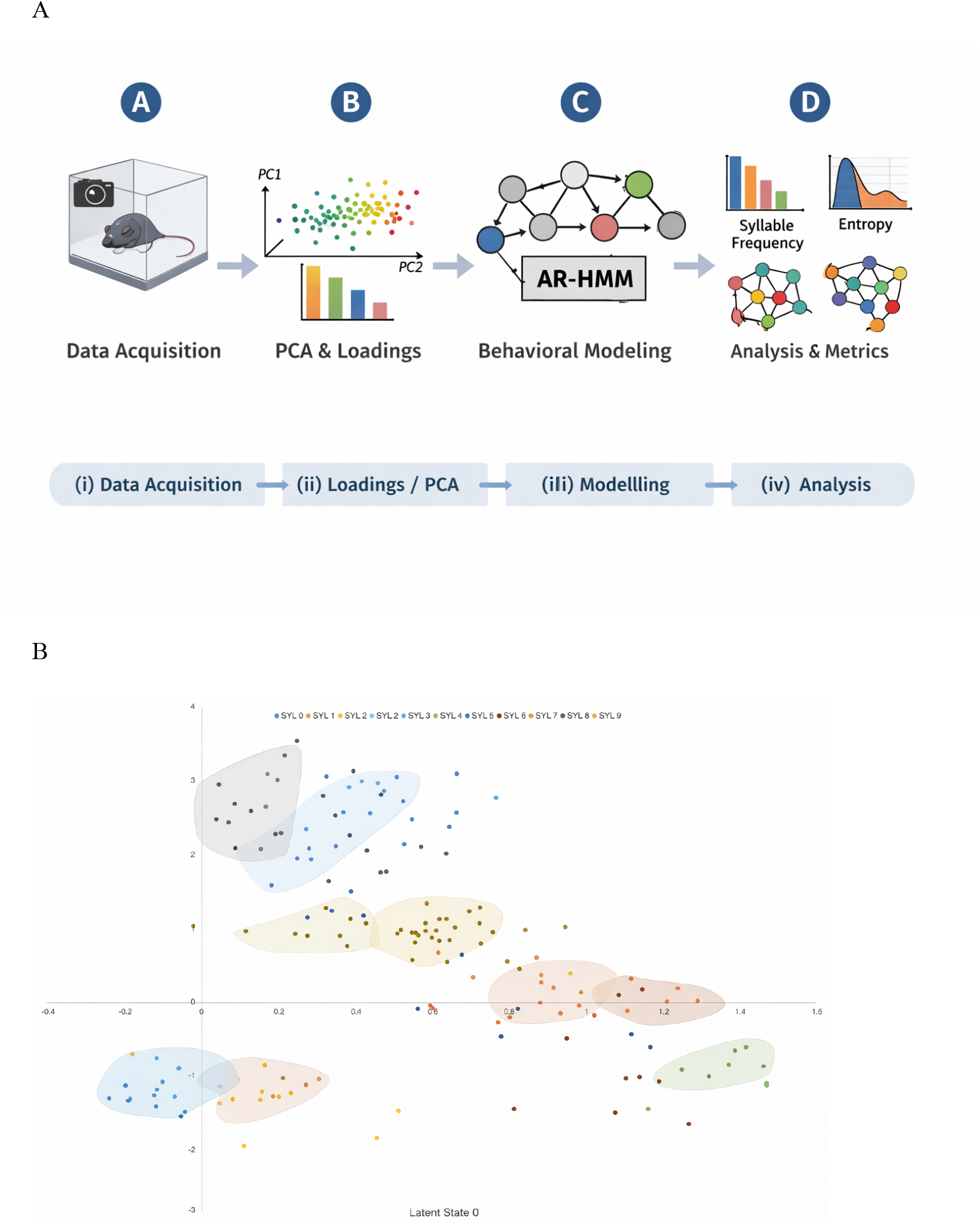
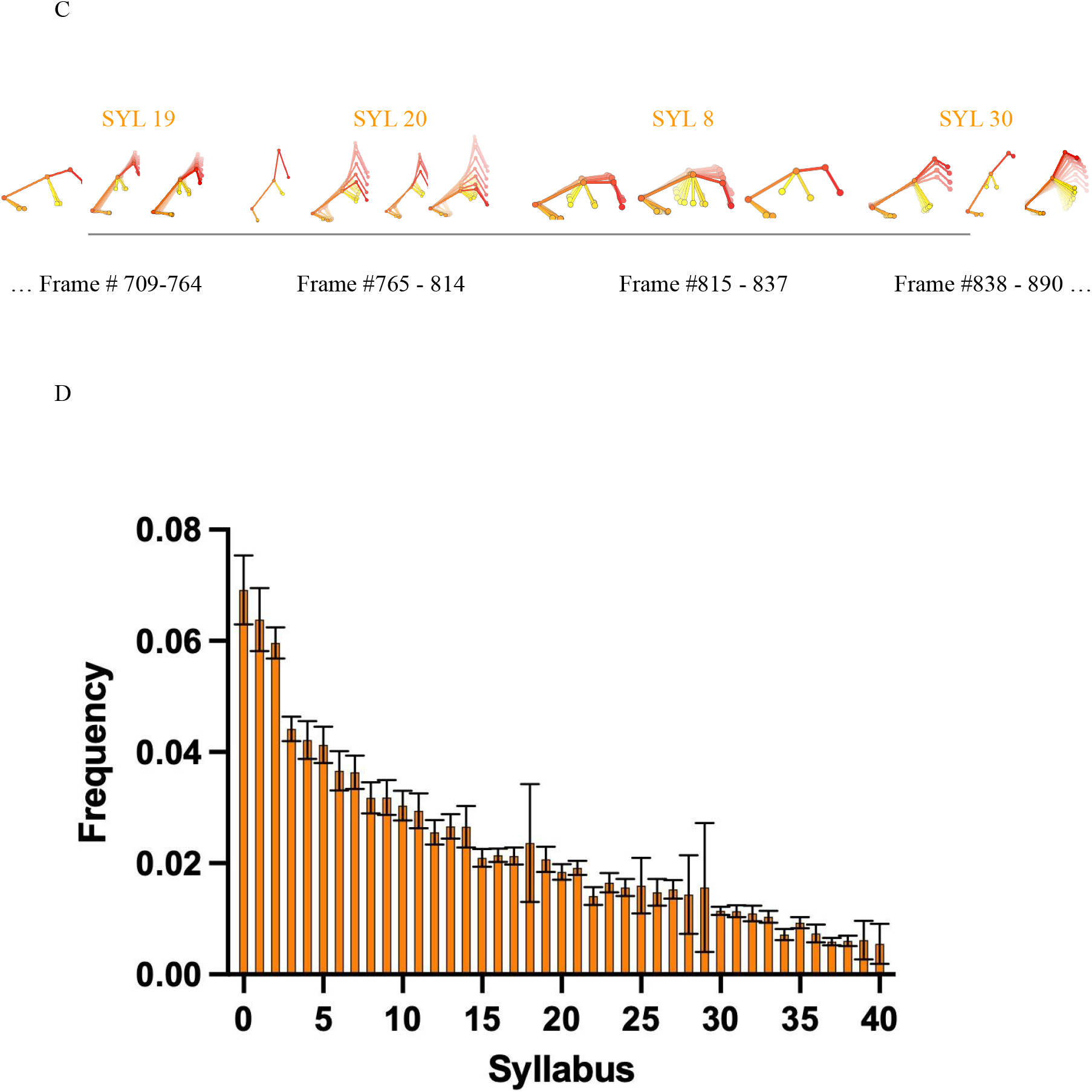
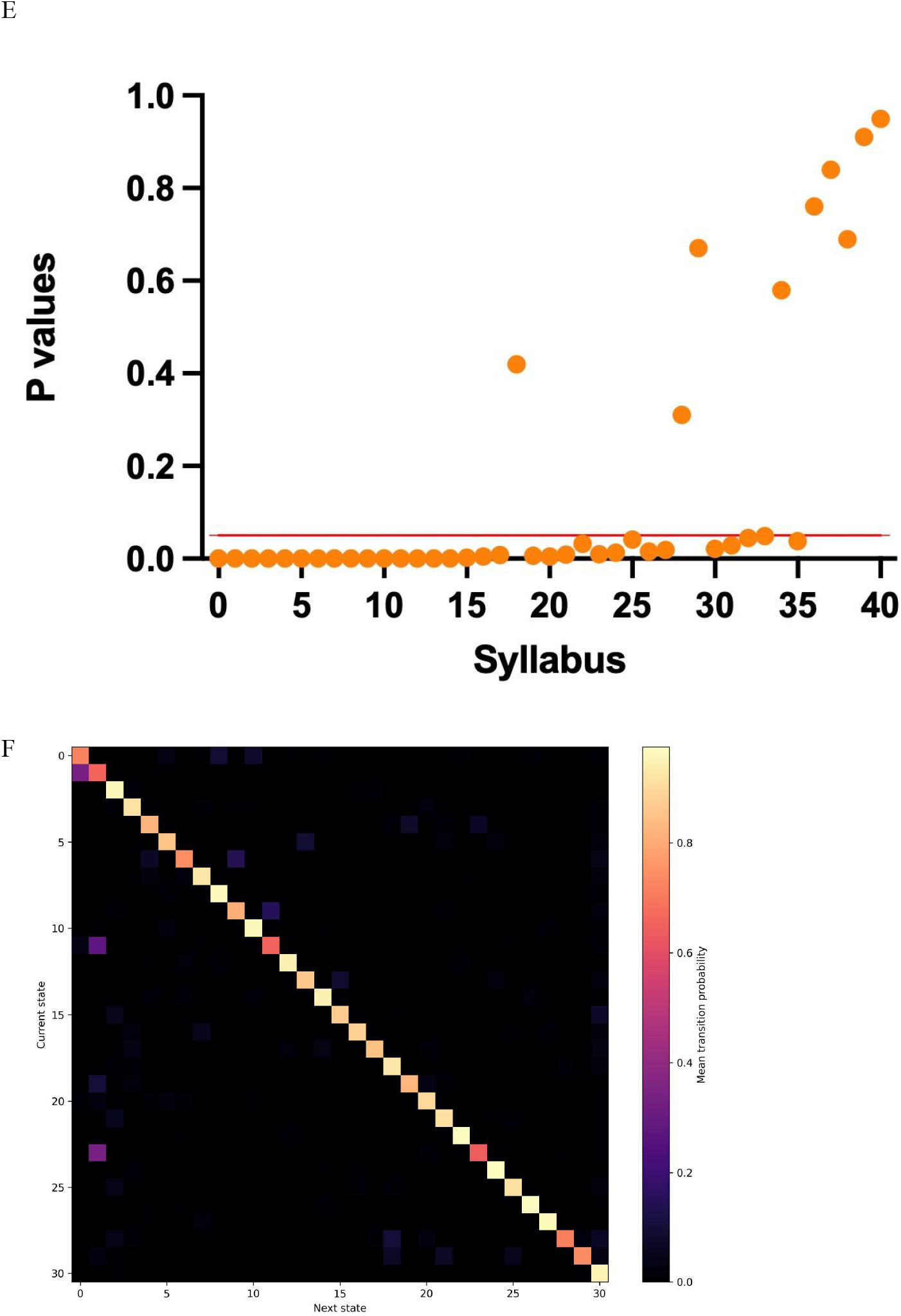
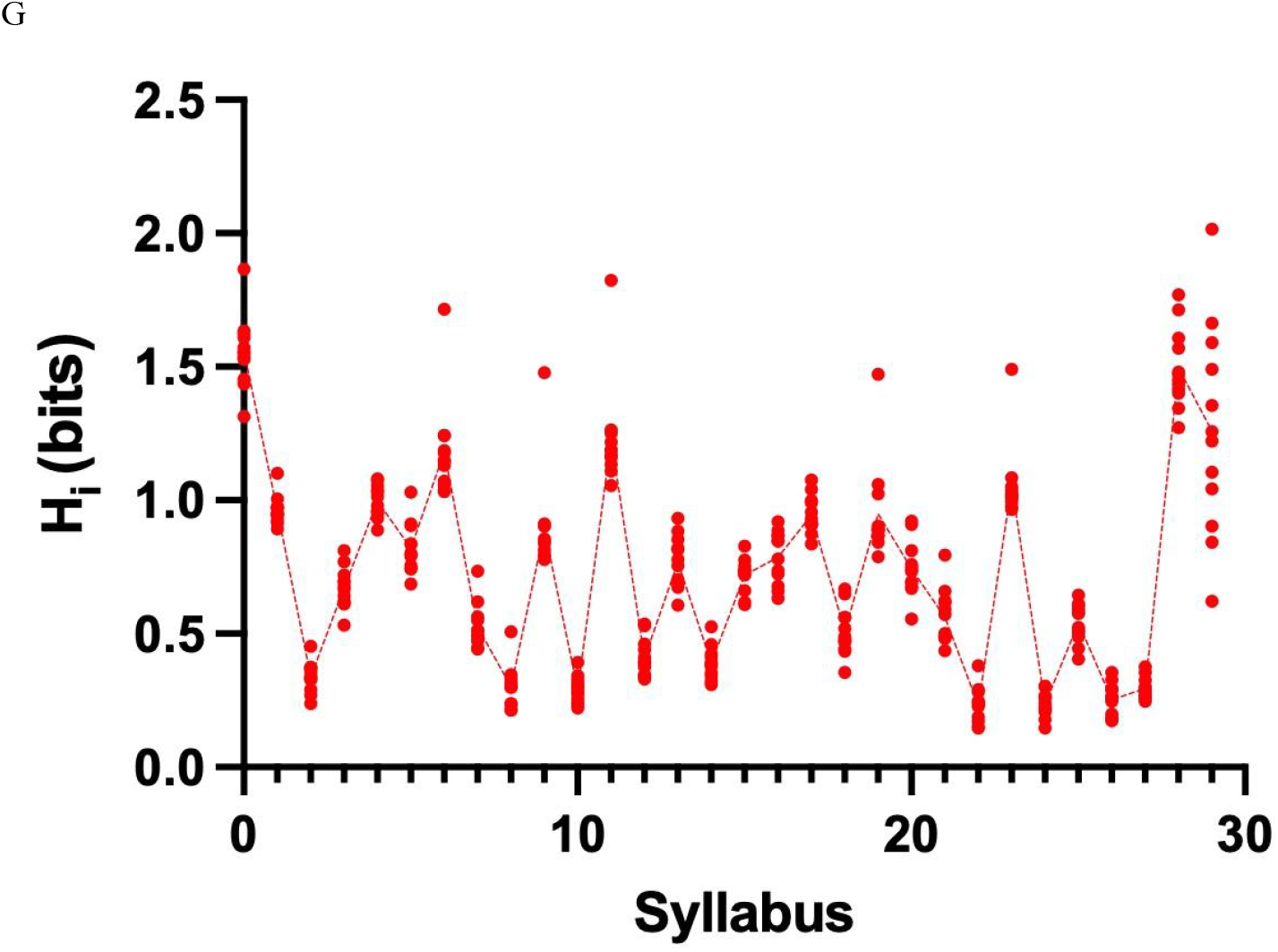
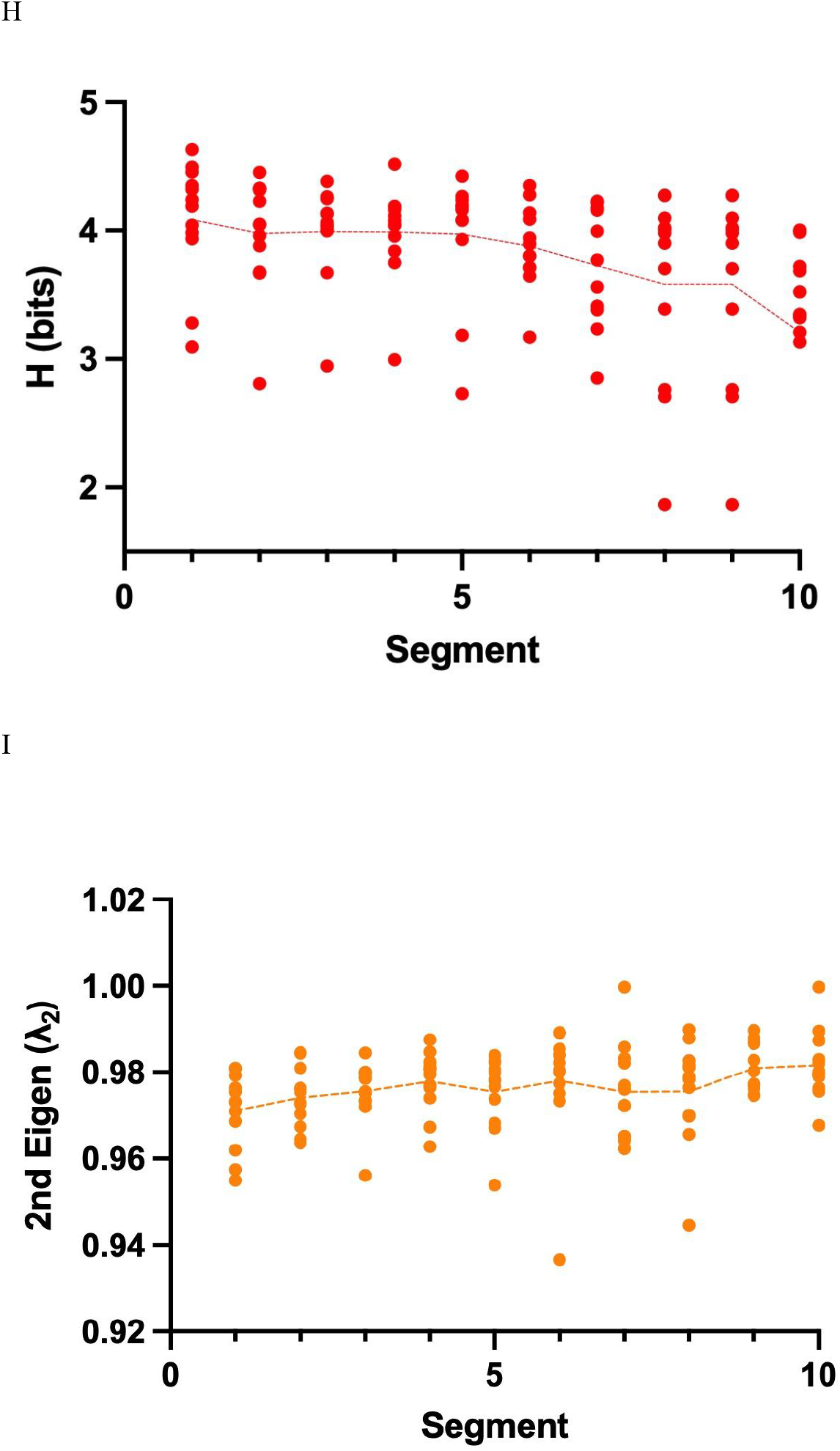
(A) Overview of the behavioral analysis pipeline. Freely moving mice are recorded using multi-camera 3D pose estimation (AVATAR-3D), keypoint trajectories are reduced via PCA, segmented into discrete behavioral syllables using an AR-HMM framework (Keypoint-MoSeq), and analyzed using frequency, transition, entropy, and spectral metrics. (B) Latent space clustering of syllables 0–9 projected onto Latent State 0 and Latent State 2. Individual frames cluster by syllable identity, demonstrating separable behavioral structure in reduced-dimensional space. (C) Example frame-by-frame syllable assignment (frames 709–890) illustrating a structured transition from syllable 19 to 20 to 8 to 30, highlighting temporally ordered behavioral segmentation. (D) Normalized syllable frequency distribution (0–40) per recording, showing structured and non-uniform behavioral repertoire usage. (E) One-sample t-test p-values assessing deviation of syllable frequencies from uniform distribution, demonstrating statistically structured state occupancy. (F) Heatmap of the cohort-averaged 31×31 transition probability matrix. Diagonal elements reflect persistence; off-diagonal structure indicates preferred state transitions. (G) Local transition entropy (Hi) per syllable (0–29 and 30+) across recordings, quantifying dispersion of outgoing transition probabilities for each behavioral state. (H) Global stationary entropy computed across 10 equal-duration segments per recording, reflecting temporal evolution of behavioral repertoire dispersion. (I) Second largest eigenvalue (Eigen_2_) of the transition matrix computed per segment, capturing temporal persistence and mixing dynamics of behavioral sequences.

An insight into the geometric and temporal structures of inferred behavioral states are represented in Figures 1B and 1C. For figure 1B, the first instances of the 10 most frequent syllables (0-9) were represented in 2 dimensional space after PCA analysis by their first two principal components, latent state 0 and 1. These clusters, color-sorted by syllables as per legend, reveals that syllables occupy separable and compact regions of the reduce-dimensional manifold; despite being crude, compared to the actual complexity of the space, the 2-dimensional simplification aimed to reveal that syllables tended to form in distinguishable hubs. The segmentation process of AR-HMM produces states that are not randomly intermingled but consistent to pose-dynamic patterns. Additionally, this separability is accompanied by a temporal component. As shown in a representative example snippet from one of our recordings (figure 1C), behavioral activity unfolds as a structured sequence of syllables, which identifiable transitions between states (e.g., syllable 19 → 20 → 8 → 30) over consecutive frames.

The structured nature of the behavior repertoire is further reflected in figure 1D. First, syllables were ranked and sorted by frequency, and the normalized occupancy probability for the syllables 0 - 40 were plotted to reveal a markedly non-uniform distribution across recordings, with certain syllables consistently expressing higher and lower frequencies than others. To formally validate that such pattern derived from random or uniform usage, a one-sample t-test for each syllable across recordings relative to the null hypothesis of equal occupancy was performed. The resulting p-values (figure 1E) demonstrated that the majority of syllables exhibit statistically significant deviations from a uniform one. In other words, differences in syllable frequencies are not attributable to sampling noise but instead reflect structured and preferential state usage. These findings confirm that spontaneous behavior in WT mice is organized around a stable, non-random repertoire of behavioral states.

The syllable annotated results from KPMS modelling and segmentation was organized into a conditional Probability Matrix P that showed the frame-by-frame transition probability upon incoming and outgoing syllable. These figures were averaged and shown as a heat map (Figure 1F). Importantly, P explicitly preserves self-transitions, allowing syllable dwell time to be incorporated into following analysis rather than collapsing each occurrence into a singular event. Hence, the diagonals are highly pronounced, with the self-transition probability being positively proportional to the average duration for each syllable. By retaining these self-transitions, the matrix captures not only which states follow one another, but also how long behavioral states tend to persist, providing a more faithful representation of spontaneous behavioral dynamics.

To further characterize syllable-specific transition structure, we computed the local entropy (Hi) for each syllable based on P (figure 1G). For each syllable, Hi quantifies the dispersion of possible next states, thereby giving it a measure of how predictable or stable transitions are, given a present state. Low local entropy indicates that a syllable transitions to a restricted subset of states, reflecting structured or stereotyped sequencing. In contrast, high local entropy reflects more distributed outgoing transitions, consistent with flexible or less constrained behavioral progression. Across recordings, syllables exhibited heterogenous Hi profiles, indicating that different behavioral states vary in degree in constraint for subsequent behavior. A general pattern can be observed, however, telling that even across different WT subjects, the degree of stability tended to be similar for each syllable, and that there exists a level of conservation in the behavioral blueprint across individuals.

Finally, we examined how global dynamical structure evolves over time by segmenting each 30-minute recording into ten equal-duration intervals and computing both global entropy (H) and spectral persistence metrics per segment (Figures 1H and 1I). Global entropy was calculated from the stationary distribution of each segment’s transition matrix, providing a measure of overall behavioral repertoire dispersion. Global entropy reflects the overall dispersion of behavioral state occupancy and complements local entropy, which quantifies transition dispersion conditional on individual states. Higher entropy values reflect broader distribution of behavioral occupancy across states, whereas lower values indicate increased constraint within a subset of syllables. Complementing this measure, the second largest eigenvalue (Eigen_2_) of the transition matrix was computed to quantify temporal persistence and mixing dynamics. Because the largest eigenvalue of a row-stochastic matrix is unity, the magnitude of Eigen_2_ captures how rapidly behavioral sequences mix over time; values closer to one indicate slower mixing and greater persistence within restricted subsets of states.

Across WT recordings, a general downward trend in global entropy and a corresponding upward trend in Eigen_2_ were observed from the first to the tenth segment. Two-tailed t-tests comparing segment 1 and segment 10 confirmed that both changes were statistically significant. Together, these shifts indicate a progressive reduction in behavioral dispersion accompanied by increased temporal persistence over the course of the session. This pattern is consistent with habituation dynamics, in which exploratory and highly variable early-session behavior gradually stabilizes into a more constrained and predictable repertoire as animals become familiar with the environment. This progressive stabilization is consistent with established principles of habituation, in which behavioral variability decreases with environmental familiarity (8). Thus, the dynamical metrics capture not only static differences in behavioral organization, but also the time-dependent stabilization of spontaneous behavior within a novel arena.

### Quantifying Pharmacological Behavioral Shifts: Spectral and Entropic Signatures of Ketamine-Induced State Reorganization

To evaluate our framework’s sensitivity to pharmacologically induced differences of spontaneous behavior, we analyzed 30-minute openfield recordings of mice administered saline or ketamine at 30 mg/kg, i.p. Six subjects were recorded for each condition. At this sub-anesthetic dose, ketamine induces NMDA receptor hypofunction and is widely used as a pharmacological model relevant to schizophrenia-associated behavioral dysregulation (9–12).

All recordings underwent identical procedures as before. Two principal findings were 1. Ketamine injected subjects sustained significantly higher levels of global entropy relative to saline controls and 2. an inversion in local transition entropy took place between ketamine-and saline favoured syllables.

Higher global entropy in ketamine-treated animals reflects a more evenly distributed behavioral repertoire, indicating that spontaneous behavior was less concentrated within a restricted subset of syllables and instead dispersed more broadly across available states. And, the inversion in local transition entropy between ketamine- and saline-favored syllables showed increasing transitional predictability and motif stabilization within some states. This dissociation suggests that ketamine simultaneously expands global repertoire dispersion while increasing local transition rigidity within specific behavioral motifs.

The syllable frequency across the two conditions were significantly different in most cases. In fact, the differential frequency analysis shown in Figure 2A showed that among the syllables 0-40, only syllables 2, 7, 11, 14, and 32-40 exhibited non-significant differences. This broad pattern indicates that ketamine does not selectively affect a narrow subset of behaviors, but rather reorganizes the structure of behavioral expression across the repertoire. In Figure 2B, the syllable differences were ranked by magnitude to explicitly display the direction and highlight syllables that were most strongly ketamine- or saline-favored.

**Figure 2.**
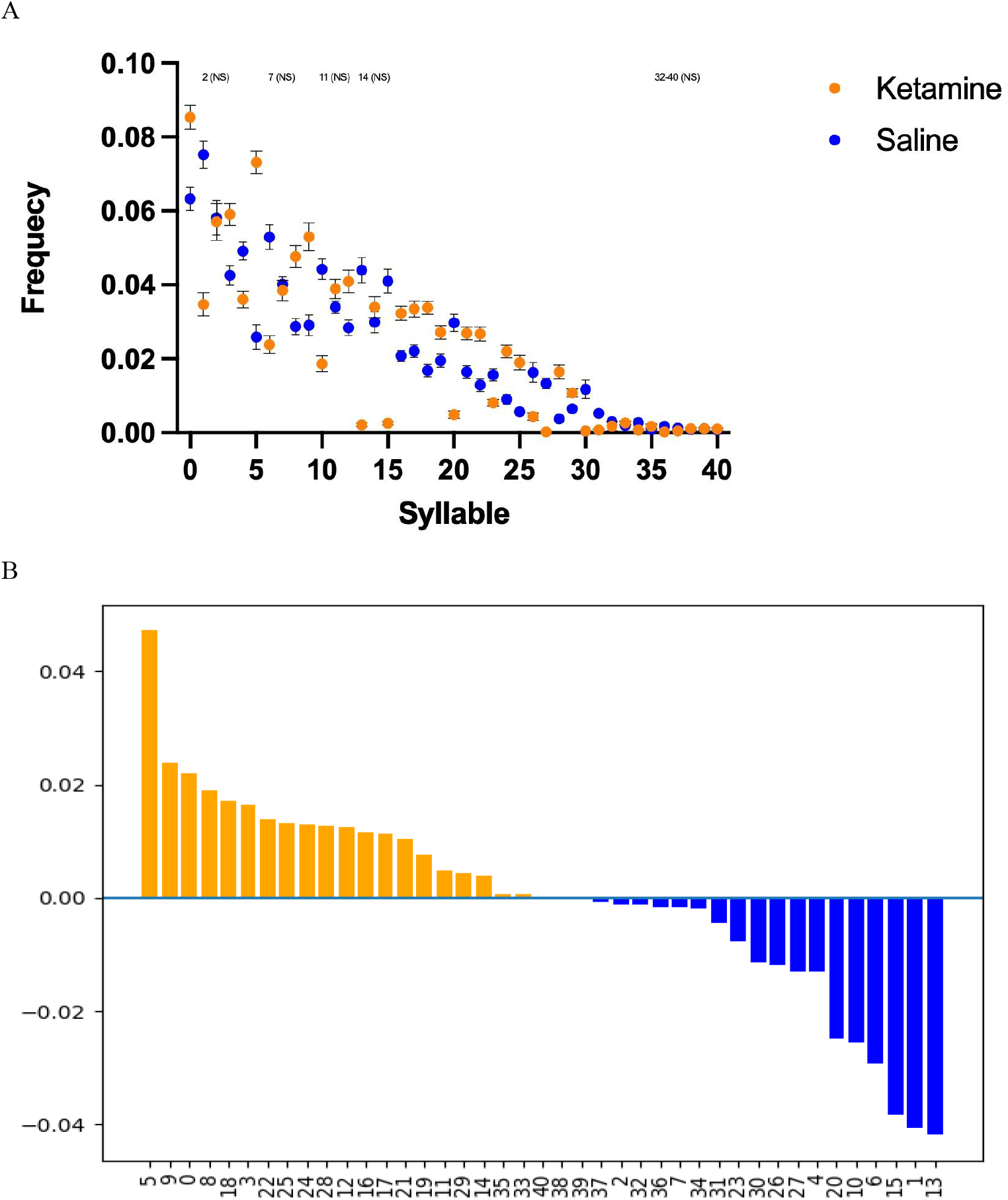
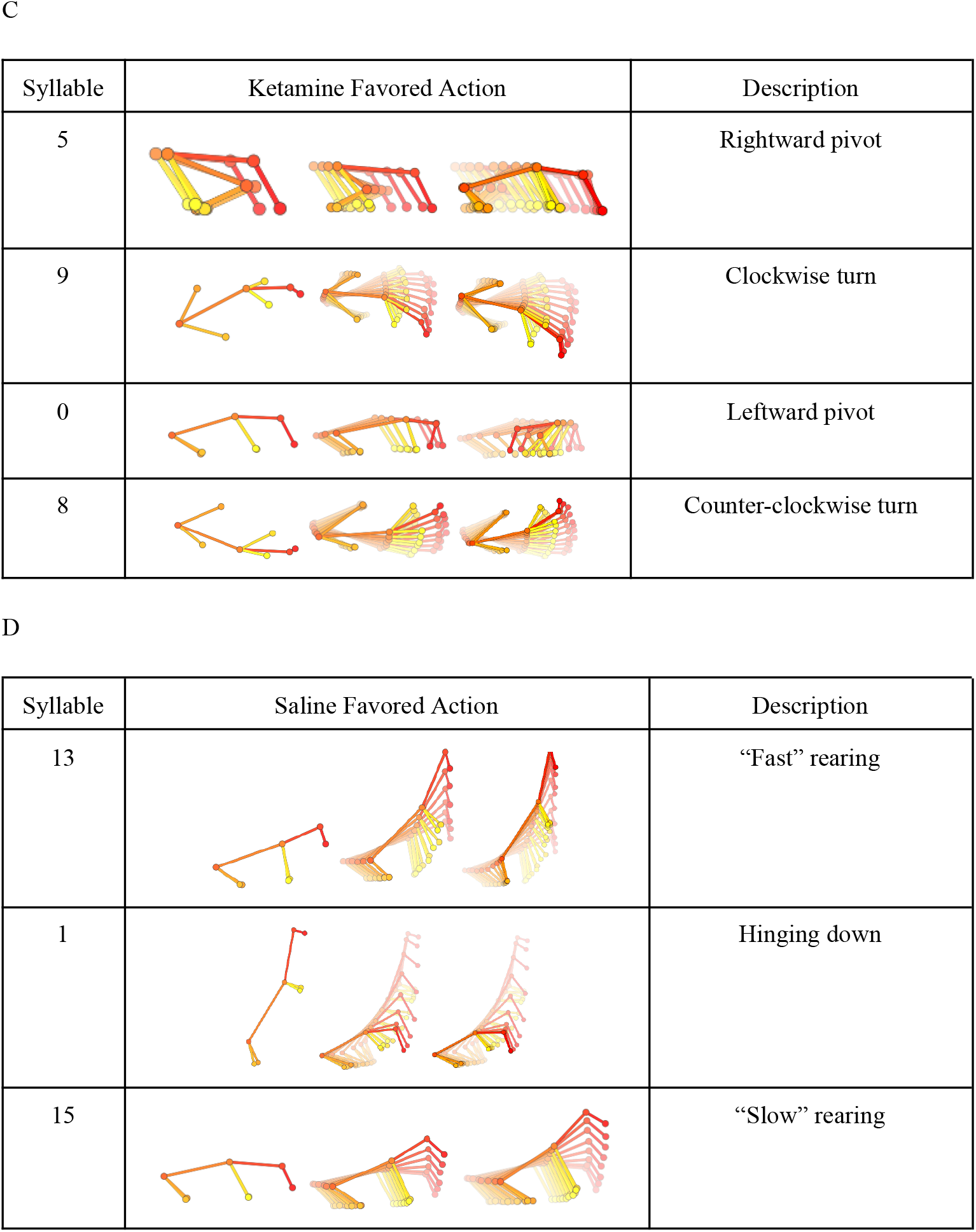
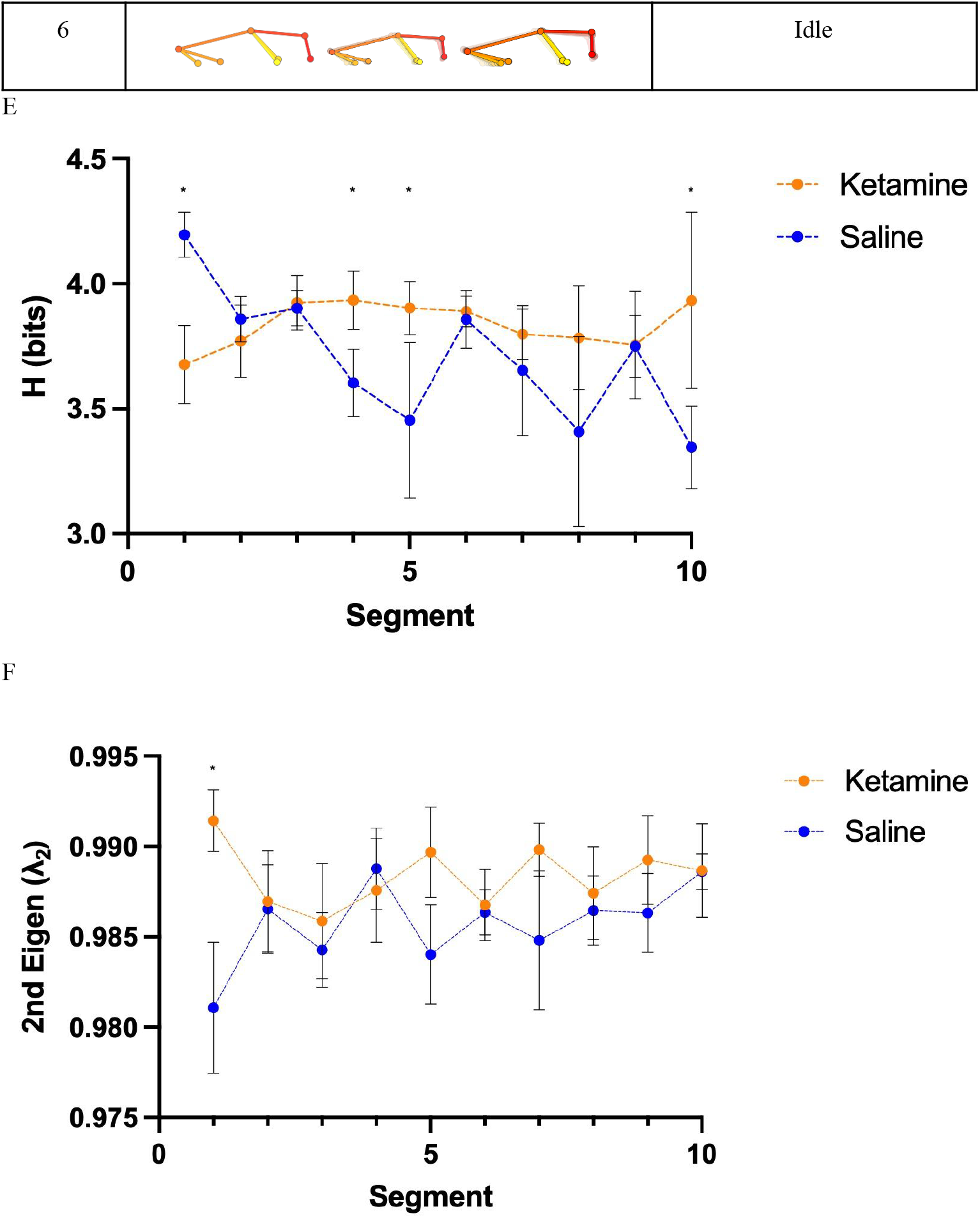
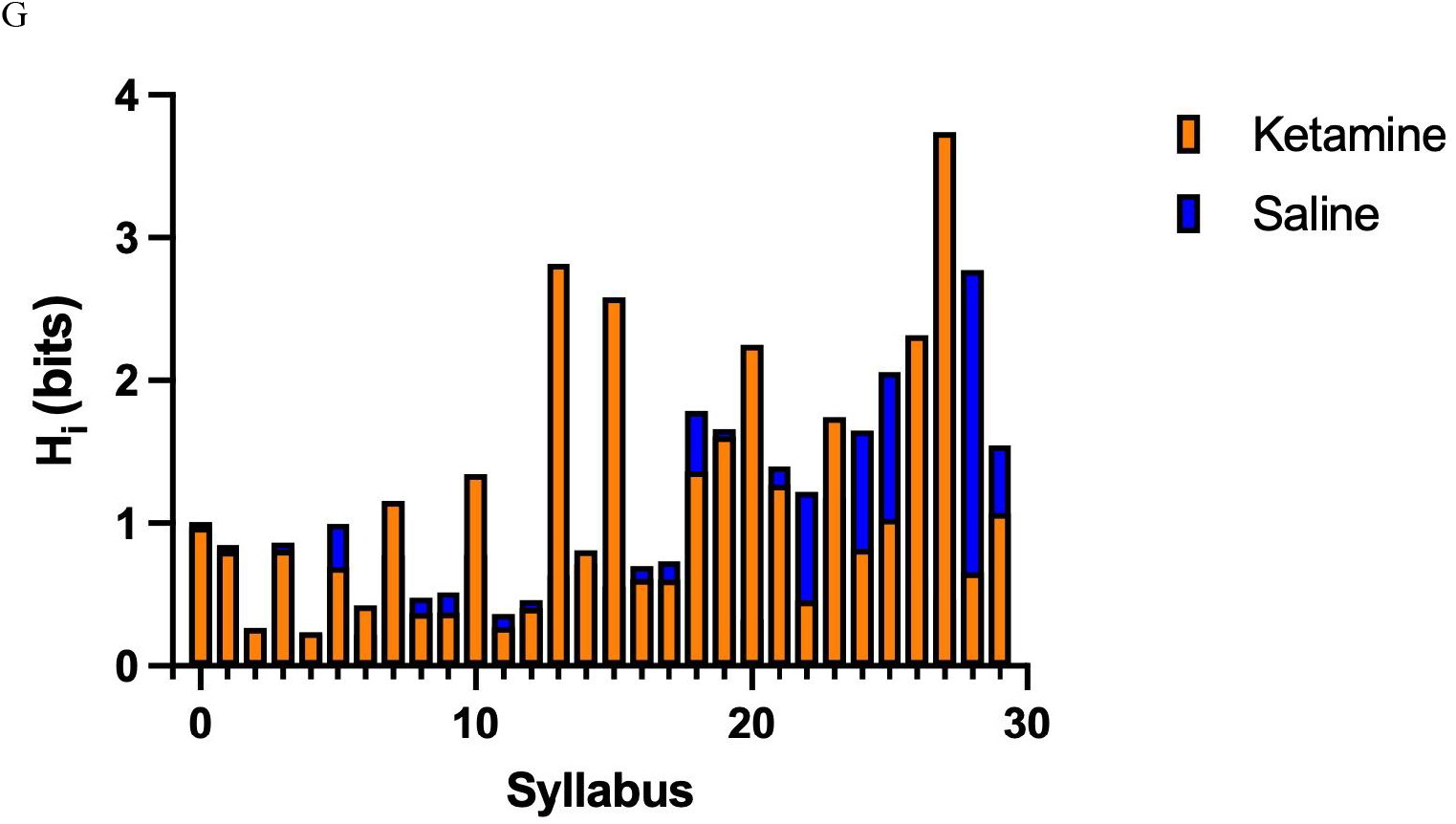
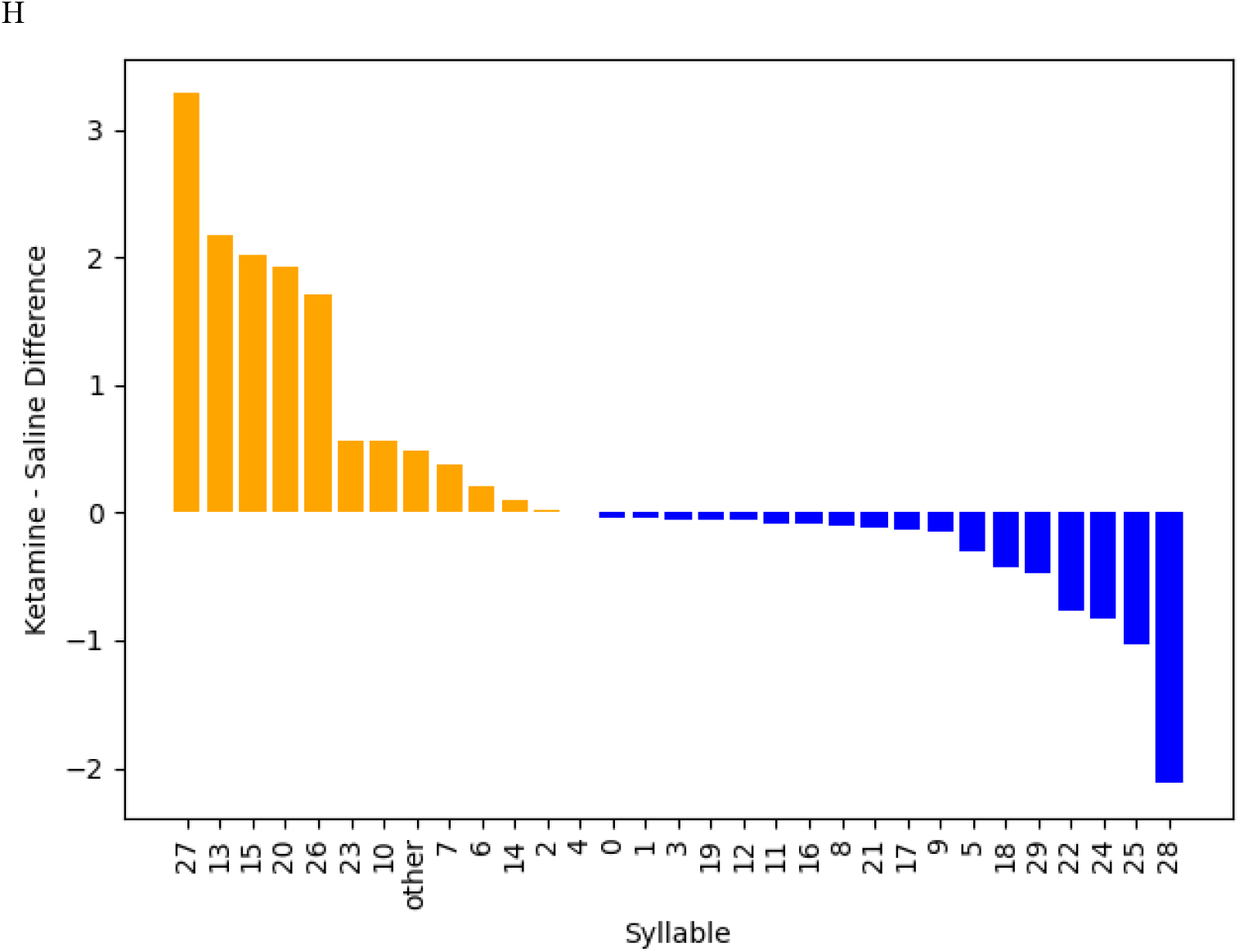
(A) Differential normalized syllable frequency (Ketamine − Saline) across syllables 0–40. Positive values indicate syllables enriched under ketamine, whereas negative values indicate syllables enriched under saline. (B) Sorted tower plot of syllable frequency differences ordered by magnitude (Ketamine − Saline). Orange bars denote ketamine-favored syllables; blue bars denote saline-favored syllables. (C) Representative frames extracted from the four most ketamine-enriched syllables, illustrating increased circling, turning, and pivoting motifs characteristic of ketamine exposure. (D) Representative frames extracted from the four most saline-enriched syllables, illustrating increased rearing, hunching, and idle behaviors characteristic of saline-treated controls. (E) Segmented global entropy (H) across time for ketamine and saline conditions, reflecting differences in overall behavioral repertoire dispersion. (F) Segmented second-largest eigenvalue (Eigen_2_) across time for ketamine and saline conditions, quantifying differences in temporal persistence and mixing dynamics. (G) Local transition entropy (Hi) per syllable under ketamine and saline conditions, measuring dispersion of outgoing transition probabilities for each behavioral state. (H) Sorted tower plot of local entropy differences (Ketamine − Saline), highlighting syllables with the greatest condition-dependent alterations in transition structure.

To characterize the types of behaviors that were most strongly ketamine or saline favored, representative frames from the four most strongly skewed syllables were visually organized (Figures 2C-2D). Ketamine-favored syllables showed increased pivoting, turning and circling whereas saline-favored syllables more prominently included reading, hinging down and idle postural behaviors. The repetitive turning motif of ketamine favored syllables aligned with stereotypical locomotive patterns for the drug. Similarly, saline favored syllables showed types of patterns associated with exploration and posture-maintenance. Together, these findings demonstrate that ketamine induces a systematic reweighting of behavioral motifs, biasing expression toward repetitive locomotor fragments while reducing engagement in exploratory and postural behaviors.

To see whether behavioral dynamics evolve over time under pharmacological effects, each recording was segmented into ten equal intervals. Global Entropy (H) and Eigen_2_ were computed and plotted for each segment (Figures 2E-2F).

While spectral analysis revealed minimal differences in Eigen_2_ between ketamine and saline treated animals across time binds, the first segment displayed significantly higher Eigen_2_ relative to saline. Given Eigen_2_ serves as a reflection of temporal persistence, this early stage increase suggests heightened short-term structural rigidity immediately following drug administration.

On the other hand, H values exhibited temporally specific differences. Ketamine-treated animals showed significantly higher entropy values in segments 4, 5, and 10. Saline treated animals saw elevated entropy in the first segment. The initially larger H value of saline treated animals may be due to heightened exploratory tendency after being introduced into the arena compounded by the proximity of injection handling. Conversely, the sustained increase in H under ketamine during later time bins suggests relatively looser and less stable behavioral structure once pharmacological effects stabilized.

Together, these findings indicate that ketamine-induced behavioral reorganization is not static but evolves across the recording session. Early post-injection dynamics may transiently alter persistence, whereas later segments reveal broader redistribution of behavioral state occupancy consistent with altered behavioral organization under NMDA receptor antagonism.

The local entropy (Hi) was calculated for both groups and its trends were compared to the frequency trend. Whereas H captures overall global repertoire dispersion, Hi gives insight into diversity of outgoing transition as well as self-transition probability from a specific state.

A pronounced inversion emerged (figured 2G-2H). Syllables that were saline favored in terms of frequency (i.e. syllables 13, 15, and 6) exhibited significantly lower higher Hi for ketamine injected subjects. Similarly, ketamine favored syllables – for instance 5, 9, 0, and 8 – showed higher Hi for saline injected subjects. Syllables that increase in frequency simultaneously become more transitionally constrained within that same condition.

This inversion suggests a multi-scale reorganization of behavioral structure. Although ketamine increased global entropy—reflecting broader redistribution of state occupancy—it reduced local entropy within ketamine-dominant motifs, indicating stabilization and increased predictability of transitions once those motifs were engaged. In parallel, saline-favored exploratory and postural states became more transitionally structured under saline. This sort of inversion between frequency and Hi sorted syllables was unique only to our ketamine and saline experiment.

## Notes

### Competing Interest Statement

The authors have declared no competing interest.

### Summary of Updates

Titles of subsections have been reorganized. Corresponding author changed to Dr. Soonwook Choi.

